# Genomic insights into metabolic flux in ruby-throated hummingbirds

**DOI:** 10.1101/2022.03.21.485221

**Authors:** Ariel Gershman, Quinn Hauck, Morag Dick, Jerrica M. Jamison, Michael Tassia, Xabier Agirrezabala, Saad Muhammad, Raafay Ali, Rachael E. Workman, Mikel Valle, G. William Wong, Kenneth C. Welch, Winston Timp

**Affiliations:** Department of Biomedical Engineering, Johns Hopkins University, Baltimore, MD, 21218, USA; Department of Molecular Biology and Genetics, Johns Hopkins University, Baltimore, MD, 21287, USA; Cell & Systems Biology, University of Toronto, 25 Harbord Street, Toronto, ON, M5S 3G5, Canada; Department of Biological Sciences, University of Toronto Scarborough, 1265 Military Trail, Toronto, ON, M1C 1A4, Canada; Department of Biology, Johns Hopkins University, Baltimore, Maryland 21218, USA; CIC bioGUNE, Basque Research and Technology Alliance (BRTA), Derio, Spain; Department of Physiology and Center for Metabolism and Obesity Research, School of Medicine, The Johns Hopkins University, Baltimore, MD 21205, USA

## Abstract

Hummingbirds are very well adapted to sustain efficient and rapid metabolic shifts. They oxidize ingested nectar to directly fuel flight when foraging but have to switch to oxidizing stored lipids derived from ingested sugars during the night or long-distance migratory flights. Understanding how this organism moderates energy turnover is hampered by a lack of information regarding how relevant enzymes differ in sequence, expression, and regulation. To explore these questions, we generated a chromosome level *de novo* genome assembly of the ruby-throated hummingbird (*A. colubris*) using a combination of long and short read sequencing and scaffolding using other existing assemblies. We then used hybrid long and short-read RNA-sequencing for a comprehensive transcriptome assembly and annotation. Our genomic and transcriptomic data found positive selection of key metabolic genes in nectivorous avian species and a deletion of critical genes (GLUT4, GCK) involved in glucostasis in other vertebrates. We found expression of fructose-specific GLUT5 putatively in place of insulin-sensitive GLUT4, with predicted protein models suggesting affinity for both fructose and glucose. Alternative isoforms may even act to sequester fructose to preclude limitations from transport in metabolism. Finally, we identified differentially expressed genes from fasted and fed hummingbirds suggesting key pathways for the rapid metabolic switch hummingbirds undergo.

## INTRODUCTION

The ruby-throated hummingbird (*Archilochus colubris*) is distinguished by features of natural and evolutionary history, morphology, and physiology from mammalian model systems such as mice, rats, and humans. They are among the smallest vertebrate endotherms (2.5-3.5 g). They employ hovering flight, displaying the highest wingbeat frequencies of any bird (and highest limb oscillation frequencies of any vertebrate; ~50-60 Hz), and in doing so sustain the highest metabolic rates among all vertebrates (R. K. Suarez 1992). In addition, ruby-throated hummingbirds engage in an annual migratory journey from breeding grounds throughout Eastern North America to wintering grounds as far south as Central America. If measured in terms of body lengths traveled, small North American hummingbirds engage in some of the longest distance aerial migrations of any species (Gass 1979). In doing so, they demonstrate a remarkable ability to sustain high rates of metabolism using endogenous lipids, an ability not shared by mice, rats, or humans (Marshall D. McCue and Pollock 2013).

To fuel these activities, hummingbirds oxidize fatty acids and carbohydrates in their flight muscles at rates faster than any other vertebrates thus far studied (Welch and Chen 2014). Remarkably, the dietary source of both fuels, carbohydrate and fat, is the same: simple sugars (glucose, fructose, sucrose) in floral nectar that provide more than 90% of the total calories they ingest (Baker, Baker, and Hodges 1998). Once ingested, hummingbirds must either oxidize or convert them into energy dense (and thus easier to carry) lipid depots. Remarkably, hummingbirds can switch between relying exclusively on oxidation of endogenous lipids to exclusive reliance on newly ingested sugars to fuel hovering flight over a period as short as 20-30 minutes (Welch and Suarez 2007; Chen and Welch 2014; Welch et al. 2006).

In order to keep up with the high energetic demands of hovering flight, hummingbirds transport, take up, and oxidize circulating sugars in flight muscles at rates as much as 55× greater than the maximum rates observed in any non-flying mammals (Welch and Chen 2014). Once in circulation, the flux of sugar to, and oxidation in, exercising muscle is thought to be limited principally at each of three key steps: 1) delivery from capillaries to the extracellular space, 2) transport across the fiber membrane, and 3) phosphorylation in the muscle fiber (Welch and Chen 2014; Wasserman et al. 2011; Rose and Richter 2005; Bertoldo et al. 2006). Mechanistic understanding of steps 2 and 3 are poorly understood as GLUT4, the key glucose transporter in mammals, is absent in birds and while hummingbird hexokinase activity is higher than other vertebrates, this alone cannot explain the rate of hummingbird glycolytic flux (R. K. Suarez et al. 2009).

The ability of hummingbirds to fuel hovering flight completely with fructose as a fuel raises interesting fundamental questions about the enzymatic basis for rapid sugar flux. The same three key steps that regulate glucose uptake and oxidation by muscles presumably apply to fructose as well (Welch and Chen 2014; Wasserman et al. 2011; Rose and Richter 2005). In hummingbirds, there is ample evidence that at steps 1 and 2 capacity for fructose uptake into flight muscle fibers is dramatically higher than in other vertebrates. However, the enzymatic basis for high rates of fructose phosphorylation (step 3) remains unknown.

While common among migratory birds (Guglielmo 2010; Jenni and Jenni-Eiermann 1998), the ability to fuel flight exclusively or predominantly with endogenous lipid stores is itself something that distinguishes hummingbirds from model mammalian species. Many avian species build fat stores to power long distance migratory flight using the fatty acids that are present in their diet (Guglielmo 2010). Some of these switch to or exploit seasonally-available diets that are rich in specific lipid classes (Guglielmo 2010; Pierce et al. 2005). However, hummingbirds achieve high rates of *de novo* lipogenesis on a simple sugar diet and high rates of lipid accumulation to see them through both overnight fasts and migratory flight.

For these reasons, genomic studies of the ruby-throated hummingbird are warranted and necessary for further understanding of these fine-tuned metabolic systems. Here we produce a chromosome level hybrid genome assembly of the ruby-throated hummingbird. We annotated the genome using a combination of Illumina and Oxford Nanopore cDNA sequencing from muscle and liver tissues to identify full coding sequences and multiple encoded isoforms. Finally, we performed differential expression analysis and differential alternative splicing analysis on fasted and fed birds in both muscle and liver tissues to fully characterize the mechanisms underlying high catalytic rates (high catalytic efficiency and/or high levels of enzyme expression) and control over metabolic flux. These results are crucial for understanding the hummingbirds’ exquisite control over rates of substrate metabolism and biosynthesis which could give insight into metabolic control of orthologous pathways in humans.

## RESULTS

### Chromosome level genome assembly

We generated a total of 26Gb of Oxford Nanopore data on the PromethION with a read-length N50 of 40Kb and 240Gb of Illumina Nova-seq data on the hummingbird brain **(Figure 1A and Methods).** We performed hybrid de novo assembly with MaSuRCA (Zimin et al. 2013) which resulted in 1,837 contigs with a contig N50 of 13.54Mb. The assembly was determined to contain 1.13% heterozygous sequence (**Figure S1).** Using the scaffolded assembly of a different hummingbird species, Anna’s Hummingbird (*Calypte anna*) (*Rhie et al. 2021*)(**Figure S2**), we performed reference based scaffolding with RaGOO (Alonge et al. 2019). Our final assembly of the ruby-throated hummingbird had 33 chromosomes that contained 98.1% of the total sequence **(Figure 1B, Table 1).** The total genome length was 1.1Gb with a scaffold N50 of 46Mb, a scaffold L50 of 5 chromosomes and the largest scaffold 100Mb **(Table 1).** We assessed the assembly for completeness using BUSCO for avian genomes (Marchi, Cirillo, and Mateo 2017) and determined it to be 96.6% complete **(Table S1)**.

**Figure 1.**
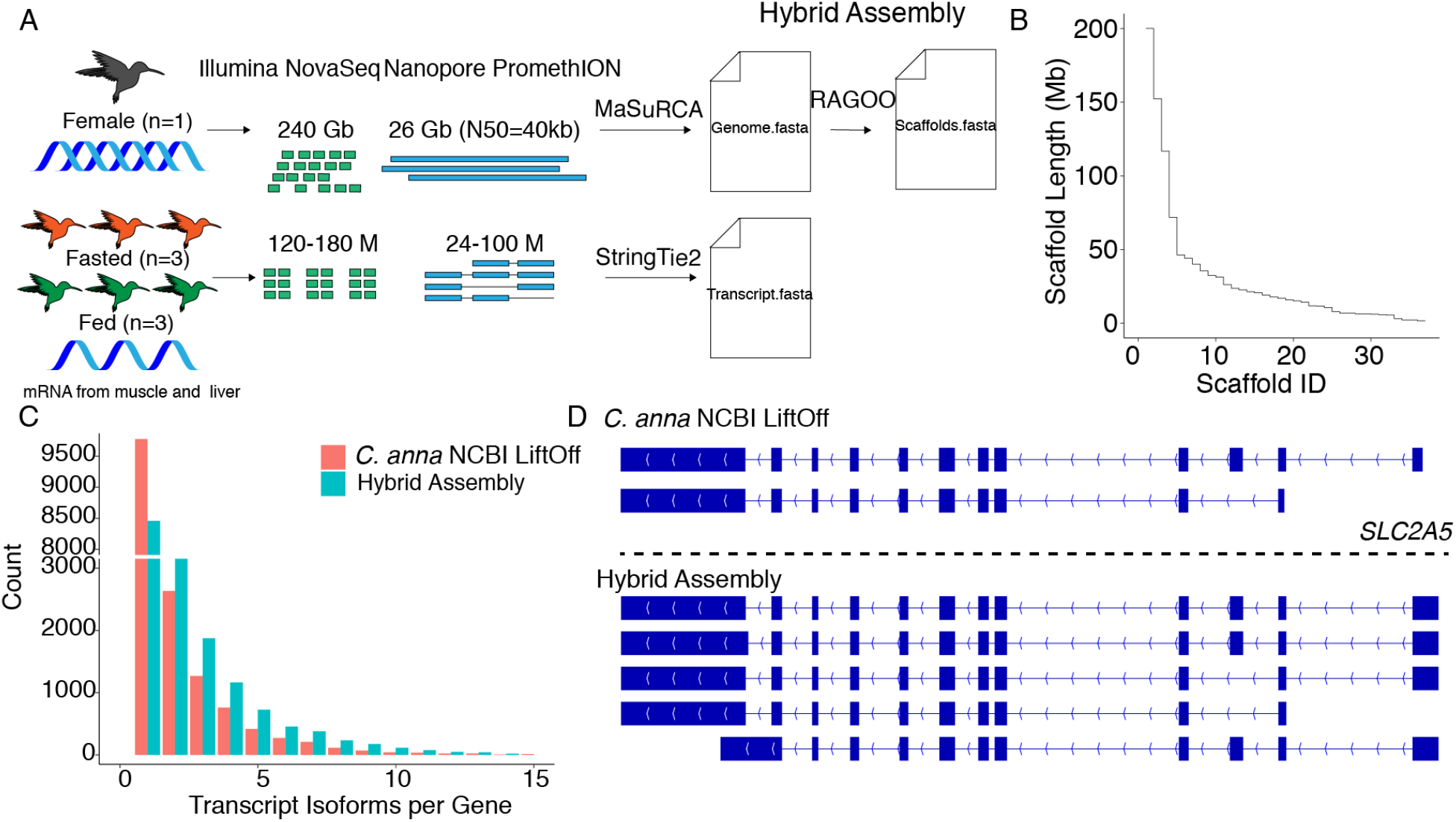
**A)** Overview of experimental design and methods for producing a chromosome level genome assembly of A. Colubris. **B)** Step plot of scaffold length for the 33 A. Colubris chromosomes. **C)** Number of isoforms per gene in the C. anna NBCI LiftOff annotation versus the Stringtie2 hybrid transcriptome assembly. **D)** The SLC2A5 gene locus in the C. anna NCBI LiftOff annotation and the Stringtie2 hybrid transcriptome assembly.

**Table 1.** Genome assembly and annotation statistics.

| *Assembly* |  |
| --- | --- |
| Total length | 1,100,571,516 |
| Chromosomes | 33 |
| Bases in chromosomes | 1,076,479,120 |
| # contigs | 572 |
| Largest contig | 200,162,936 |
| N50 | 46,260,142 |
| L50 | 5 |
| *Genomic Features* |  |
| %GC | 42.05 |
| %TE | 10.5 |
| %Repetitive | 14.83 |
| *Gene Repertoire* |  |
| Genes | 17,878 |
| Transcripts | 43,348 |

### Genome Annotation

Avian genomes are the smallest of the amniotes, with smaller genomic elements (*e.g*. introns, exons, intergenic DNA) and fewer transposable elements compared to mammals (Zhang et al. 2014). With our new genome in hand, we examined the repetitive elements in the *A. Colubris* genome assembly. We used RepeatModeler2 to generate *de novo* repeat libraries (Flynn *et al*. 2020) and used them in combination with the curated Avian library to perform homology based repeat masking with RepeatMasker. Among vertebrates, birds exhibit relatively low copy numbers and an overall reduced diversity of repetitive elements (International Chicken Genome Sequencing Consortium 2004; Dalloul et al. 2010; Warren et al. 2010; Sotero-Caio et al. 2017), with the exception of the woodpecker (*Picoides pubescens*) whose genome is 22.2% TEs, mostly contributed by the LINE/CR1 (Zhang et al. 2014). In *A. Colubris* we detected 163 Mb of repetitive sequence representing 14.83% of the genome, including 116Mb of TEs that make up 10.50% of the genome, consistent with the repeat content in other avian lineages **(Table 1, Table S2)**(Zhang et al. 2014). Among classified repeats, LINE/CR1 elements were the most abundant superfamily found in the *A. Colubris* genome, making up 6.95% of the sequence. Next were LTRs (2.57%) and repeats discovered by our *de novo* libraries but not classified by RepeatClassifier (Unknown; 2.50%). Preliminary gene annotation was accomplished via a liftover of the *C. anna* annotations from the NCBI annotation (GCF_003957555.1) with LiftOff, a tool that maps annotations between closely related species (Shumate and Salzberg 2020). The *C. anna* annotation LiftOff to *A. colubris* consisted of 15,879 genes and 31,163 transcripts for an average of two transcripts per locus.

### Transcriptome assembly

In order to capture both the complexity of differential splicing and the precision of splice junctions, transcription start and end sites we used a combination of short-read Illumina NovaSeq and long-read Oxford Nanopore cDNA sequencing on six hummingbirds across both muscle and liver tissue **(Figure 1A).** Briefly, we used the hybrid reference based assembly pipeline from StringTie2 (Shumate et al. 2021) to expand our existing *C. anna* LiftOff annotation to a total of 17,878 genes and 43,348 transcripts. Our transcriptome assembly identified 96.4% (41,807) of genes containing multiple isoforms with an average of 2.4 isoforms per gene, a large improvement over the *C. anna* NCBI LifOff annotation alone **(Figure 1C)**. Additionally, our assembly identified 1,999 novel loci of which 1,051 were functionally annotated by BLASTing to the SwissProt database. Included in these novel genes are genes critical to metabolism including *ALDOA, PFKM, G6PD, PGLS, PC, PCK2, PFKFB1, PYGM*, and *PLPPR1*. Furthermore, the hybrid transcriptome assembly increases the number of isoforms variants per gene as exemplified by the Solute Carrier Family 2 Member 5 (*SLC2A5/GLUT5*) where the *C. anna* NCBI annotation contained two transcripts and our new hybrid annotation contains five splice isoforms that encode for different protein isoforms **(Figure 1D)**. Our expanded annotation provides the opportunity to understand gene expression changes at the transcriptome level during transitions between fuel use regimes, thus providing insights into potential mechanisms that make these organisms such flexible metabolic performers.

### Positively selected genes in nectivory

Nectar feeding animals have among the highest recorded metabolic rates, incidentally, flight requires the highest metabolic rates of any form of locomotion known (Raul K. Suarez, Herrera M, and Welch 2011; R. K. Suarez 1992), with metabolic rates reaching 170 times higher than at rest (Welch and Chen 2014). Using our new ruby-throated hummingbird genome assembly, we performed phylogenetic analyses of nectivorous avian species. We used 20 species, (Chimney swift, Anna’s hummingbird, Helmeted guineafowl, Chicken/Red junglefowl, Wild turkey, Japanese quail, Zebra finch, Bengalese finch, Common canary, Painted honeyeater, Black sunbird, Cape sugarbird, Emperor penguin, Adelie penguin, Burrowing owl, Barn owl, African ostrich, Sanda bush warbler, Hooded crow, ruby-throated hummingbird) of which five are nectivorous from four separate lineages **(Figure S3)**. We used OrthoFinder (v2.3.12) to identify orthologous gene clusters between all 20 species (Emms and Kelly 2019). OrthoFinder groups genes into orthogroups, sets of genes descended from a single gene in the species, the last common ancestor based on their sequence similarity. OrthoFinder assigned 98.0% of genes to orthogroups generating 17,895 orthogroups containing a total of 364,583 genes. Of these orthogroups, 5,085 (28%) were shared between all 20 species and 1,207 were shared and present as a single copy (**Figure 2A**).

**Figure 2.**
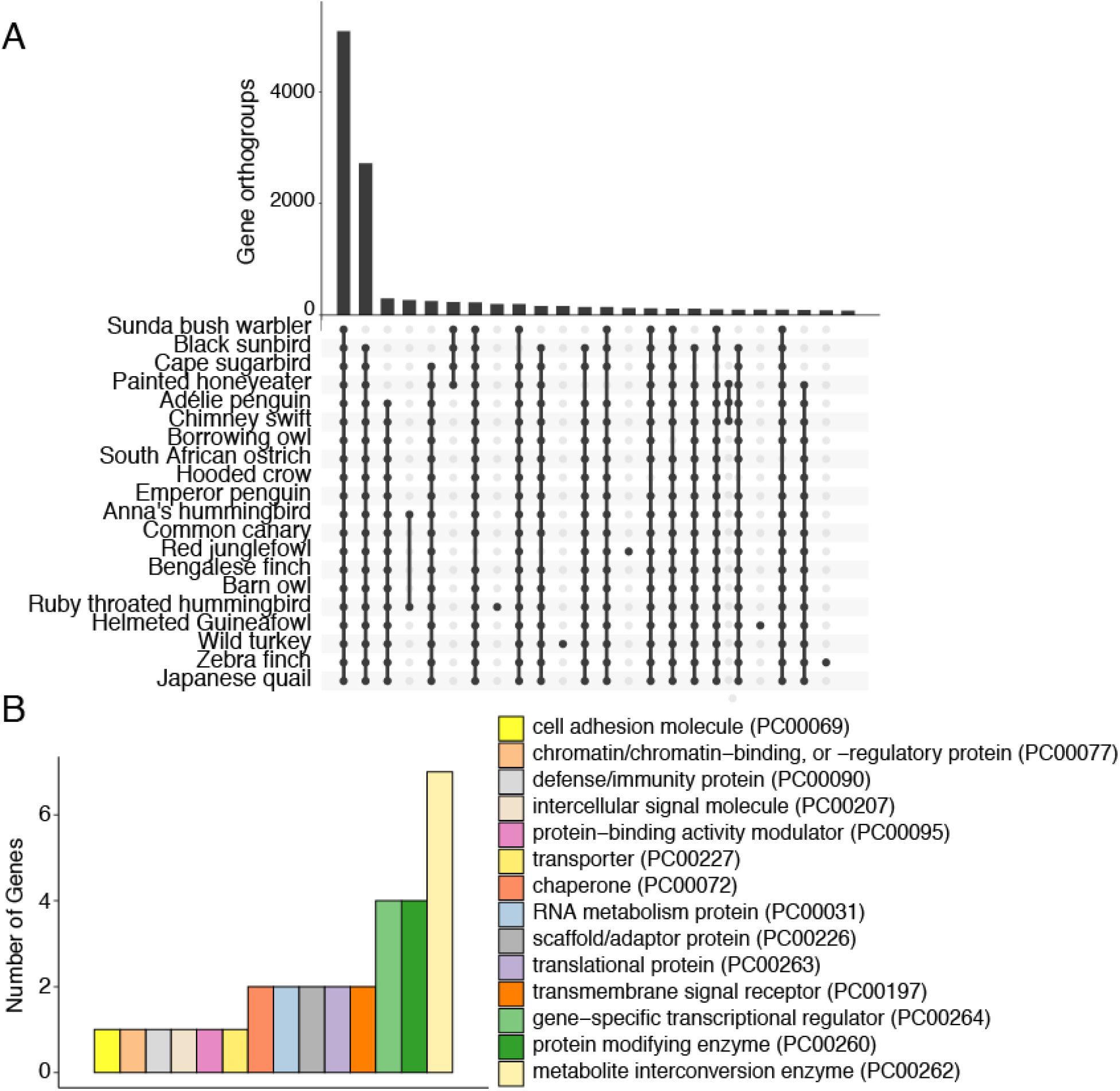
Positive selection in nectivory. **A)** Upset plot illustrating the number of shared orthogroups between the twenty species. Bars with less than 100 orthogroups were removed. **B)** Panther gene ontology classification of the positively selected genes.

Next, we analyzed all shared single copy orthologous proteins for evidence of positive selection with PAML (Yang 2007) (**Methods**). Of the 1,207 shared single-copy orthologs we determined 39 (3.2%) genes with evidence of positive selection in the nectivorous branches **(Table S3)**. A gene ontology (GO) analysis revealed that seven of the positively selected genes are metabolic interconversion enzymes **(Figure 2B)**. Nectivorous birds have extremely high rates of substrate metabolism, therefore it is likely that enzymes in these pathways are well adapted to increase the rate of sugar metabolism. We found two enzymes crucial for oxidative sugar metabolism to be positively selected, *PDHA1* and *GAPDH. GAPDH* is the enzyme separating lower and upper glycolysis and is the rate limiting step in the pathway (Shestov et al. 2014). *PDHA1* is a subunit of the pyruvate dehydrogenase complex in the mitochondrial matrix that controls the flux of pyruvate (the end product of glycolysis) into the tricarboxylic acid cycle. With an extremely low fat diet, nectivorous birds rely on rapid lipogenesis to generate endogenous fat storage and lipolysis for fueling flight in the fasted state. Interestingly, we identified positive selection in genes involved in fatty acid elongation (*ACADL*), beta-oxidation (*HACD3*) and ketone utilization (*BDH2*) (Guo et al. 2006). These data indicate that select genes (and the pathways they regulate) are positively selected in nectivorous birds to enable their highly energetically demanding lifestyle.

### Hummingbird sugar transport and metabolism

The relative expression of the distinct GLUT transporters across the liver and muscle tissues provides key insights into hummingbird sugar metabolism. In the liver tissue the primary GLUT genes are *SLC2A2* and *SLC2A5* with a medium level of transcription of *SLC2A9, SLC2A10* and *SLC2A11* and comparatively low levels of *SLC2A1, SLC2A3, SLC2A6* and *SLC2A13* **(Figure 3A).** The muscle tissue has the highest expression of *SLC2A5*, medium expression of *SLC2A1* and *SLC2A12* and low levels of *SLC2A10, SLC2A3, SLC2A11, SLC2A13* and *SLC2A2. SLC2A2*, encoding the GLUT2 protein which has a high K_m_, plays a stronger role in enteric (Karasov 2017) and hepatic (Mueckler and Thorens 2013) sugar transport, resulting in the expected higher expression we observe in liver over muscle samples. Interestingly, chicken *SLC2A1* and *SLC2A3* share sequence homologies of ~80% and ~70% respectively with human GLUTs, but other isoforms such as *SLC2A2* and *SLC2A5* only share ~65% and ~64% sequence homology (Ali et al. 2020). A comparison of *SLC2A2* sequences to 20 bird species reveals the loss of an *N*-linked glycosylation site in four of the 20 species (Workman et al. 2018). In the case of the African ostrich and the barn owl this site is lost due to truncation at the 5’ end of the protein. However, in the hummingbirds (Anna’s hummingbird and the ruby-throated hummingbird) the Asn-64 is replaced by Ser-64, therefore eliminating the conserved *N*-linked glycosylation site present in the sixteen avian species as well as humans and mice. In mice, the loss of this glycosylation site is coincident with increased GLUT2 protein endocytosis and the onset of type 2 diabetes (Ohtsubo et al. 2005).

**Figure 3.**
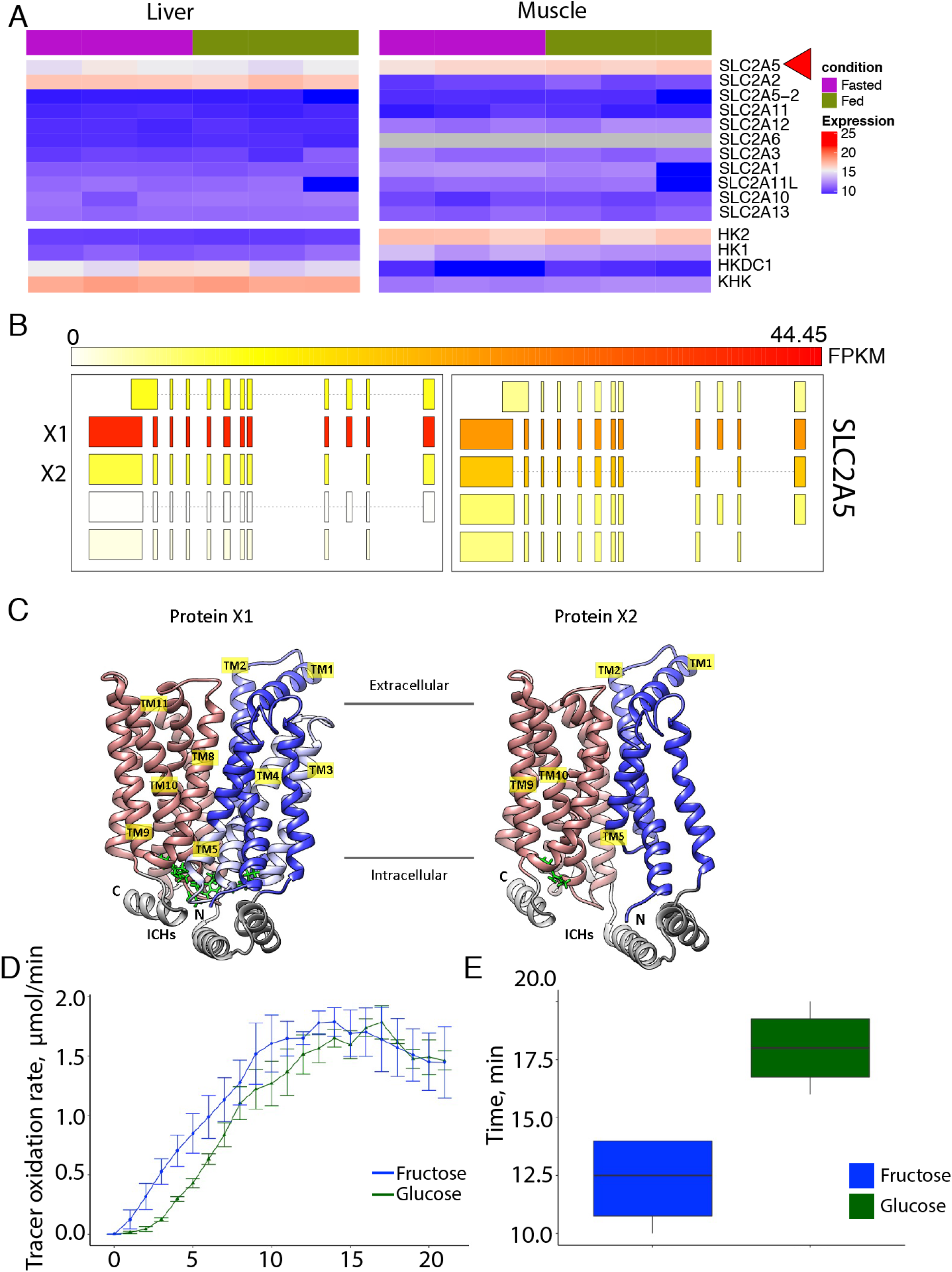
Hummingbird sugar transporters. **A)** *Gene expression heatmaps for ruby-throated hummingbird muscle and liver tissue. Variance-stabilizing transformation is applied for graphical representation. SLC2A6 is only expressed in the liver, therefore boxes in muscle are filled in as gray*. **B)** (*Left) Isoform expression of SLC2A5 in the liver and (right) muscle. FPKM of each isoform color coded according to the top scale bar*. **C)** Ribbon representations of the two *protein models for GLUT5 isoforms predicted by Alphafold2 based on the mammalian SLC2A5 ortholog. Left is the X1 isoform, right is the X2 isoform that is missing exon 3. In both atomic models amino- and C-terminal TM bundles are colored blue and red respectively. Regions of the X1 isoform that are missing in the X2 isoform are depicted in light blue. Arginine-glutamate salt bridges at the intracellular tips of TMs are green. ICHs stands for “intracellular helices”. **D)** Tracer oxidation rate over twenty minutes when birds were fed ^13^C on either the glucose or the fructose portion of the disaccharide. **E)** Peak oxidation time of glucose and fructose in minutes (p=.02, pair t-test*).

The absence of *SLC2A4* (GLUT4) leaves many unanswered questions about how glucose enters avian muscle cells. From our study we note particularly high liver and muscle expression of *SLC2A5* (GLUT5), which facilitates fructose uptake in mammals (Barone et al. 2009). *SLC2A5* is not expressed highly in mammals and in mammalian GLUT5 a single point mutation is enough to switch the substrate binding preference of GLUT5 from fructose to glucose (Nomura et al. 2015). The abundance of *SLC2A5* transcripts in hummingbird tissues, especially muscle tissue, is particularly interesting because it suggests this transporter is principally responsible for glucose/fructose transport into hummingbird tissues. There is considerable sequence divergence between hummingbird GLUT5 and mammalian GLUT5 (65.5% identity to human, 63.7% identity to mouse) and even from hummingbird to chicken (80.5% identity). We hypothesize that this form of hummingbird GLUT5 has transport capacity for glucose, but at a lower affinity than its capacity for fructose. In bacterial GLUT transporters (e.g. *XylE*) a Trp residue at the floor of the sugar binding pocket displays two hydrogen bonds with the bound glucose (Sun et al. 2012). In the same position on rat GLUT5 this residue moves to Alanine (Ala395) (Nomura et al. 2015) and in the hummingbird this residue is serine (Ser403) **(Figure S4)**. The Trp amino acid is well conserved amongst all human glucose transporters (GLUT1-4), however it moves to Ser in human GLUT7 which is also a dual (glucose and fructose) transporter **(Figure S4)**. With this information we can speculate that hummingbird GLUT5 could be a dual glucose/fructose transporter.

Using long read cDNA data we quantified the relative abundance of *SLC2A5* transcripts in muscle and liver and identified differential alternative splicing occurring between muscle and liver tissues **(Figure 3B)**. Particularly, the muscle tissue has higher expression of the isoform that skips exon 3. The dominantly transcribed isoform translates to a protein highly similar in structure to mammalian GLUT5 (Nomura et al. 2015). However, the muscle GLUT5 variant skipping exon 3 is missing transmembrane domains TM3, TM4 and the intracellular tip of TM5 **(Figure 3C).** In this isoform, the salt bridges between the amino and C-terminal TM bundles are absent and therefore the outward facing state is likely not favored. Our transcriptome sequencing in two different tissues across two opposing metabolic states (fed and fasted) highlighted the complexities of metabolic regulation at the transcriptional level.

While the high expression of fructose transporter gene *SLC2A5* strongly suggests that fructose uptake capacity may be sufficient to meet fructolytic and oxidative demand during hovering flight, the enzymatic basis for high rates of fructose phosphorylation is still unclear. The main sugar kinase expressed in the liver is ketohexokinase (*KHK*), which has high affinity for fructose in mammals. However, in both humans and in the ruby-throated hummingbirds, the muscle mainly expresses hexokinase 2 (*HK2*), which is a glucose-specific kinase in humans **(Figure 3A)**. The hummingbird *KHK* and *HK2* genes have 65% and 87% identity to their human orthologs, respectively, therefore their substrate affinities could be different from their human orthologs.

Previous studies assessed *A. Colubris* muscle total hexokinase activity and determined the V_max_ to be 50% lower for fructose than glucose phosphorylation, which would not keep up with the calculated required rates of fructose oxidation by flight muscle during hovering flight (Myrka and Welch 2018). To further understand differences in fructose and glucose metabolism we used a chronic stable isotope tracer methodology to examine the speed of glucose and fructose usage for *de novo* lipogenesis in the ruby-throated hummingbird. We fed ruby-throated hummingbirds sucrose-based diets enriched with ^13^C on either the glucose or the fructose portion of the disaccharide. Isotopic incorporation into fat stores was measured via the breath ^13^C signature while fasting (oxidizing fat). We found that the respiratory exchange ratio 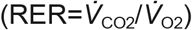 (RER) and tracer oxidation increased quickly with feeding (**Figure 3D, Figure S5A**), and with the respiratory exchange ratio RER exceeding a ratio of 1, suggesting lipid synthesis was occurring along with tracer oxidation. At the 10 min mark the RER began to fall but remained above 0.85 for the remainder of the trial. Peak tracer oxidation did not differ between the enriched sucrose solutions (**Figure S5B**, p = 0.66). However, the time to peak oxidation differed, with the fructose-enriched sucrose solution reaching peak tracer oxidation faster than the glucose-enrich sucrose solution (**Figure 3E**, p = 0.02). Overall these data support a hypothesis where fructose is rapidly transported out of the blood and metabolized while glucose remains in the bloodstream for longer and is used as a fuel source when blood fructose levels decline.

Another key regulator of blood glucose homeostasis is glucokinase (*GCK*), in mammals this enzyme has a high K_m_ and is the glucose sensor not only for regulation of insulin release by pancreatic β-cells, but also for key organs that contribute to glucose homeostasis, such as the liver (Matschinsky and Wilson 2019; Peter et al. 2011). However, birds do not express GLUT4, the insulin sensitive glucose transporter, and the ruby-throated hummingbird in particular maintains the highest blood glucose concentration known amongst vertebrates (Ali et al. 2020; Beuchat and Chong 1998). Our transcriptome assembly did not identify *GCK* in the assembled hummingbird transcriptome and we did not identify any *GCK* sequence in any of the ruby-throated hummingbird RNA-seq reads. When we compared the ruby-throated hummingbird genome to the chicken reference genome we determined that the region of the genome containing the GCK gene is not syntenic to any of the hummingbird sequence **(Figure S6)**.

### Identification of differentially expressed genes that respond to fasting

To identify differentially expressed genes (DEGs) that rapidly respond to fasting, we profiled the transcriptomes of total mRNA from the muscle and livers of *A. Colubris* hummingbirds that were fed sucrose *ad libitum* (fed) or fasted for a time period of one hour (fasted) **(Figure 1A).** We analyzed three biological replicates for each metabolic condition (fasted versus fed) with Stringtie2 hybrid long and short-read quantification and DESeq2 with our newly constructed reference and annotation (Love, Huber, and Anders 2014; Shumate et al. 2021). The fasted versus fed condition produced marked change in the transcriptomes. We identified 140 differentially expressed genes (DEGs) with adjusted p-values below 0.1 in the liver **(Figure 4A, Table S4)** and 191 DEGs in the muscle **(Figure 4C, Table S5).** Thus, the one hour fasting targets targeted a relatively small set of genes in the muscle and liver that likely play a role in the hummingbird’s rapid switch from fed to fasted metabolism. To categorize these genes according to their gene ontology we used the Genetonic pipeline and generated functional gene-set enrichments for both the *A. Colubris* liver and muscle (Marini et al. 2021) **(Figure 4B,D)**. This analysis yielded 200 statistically significant pathways (FDR < .05) in the liver and 106 in the muscle (**Table S6-7)**. The response to fasting in the muscle and liver influenced dramatically different metabolic and regulatory pathways in each tissue.

**Figure 4.**
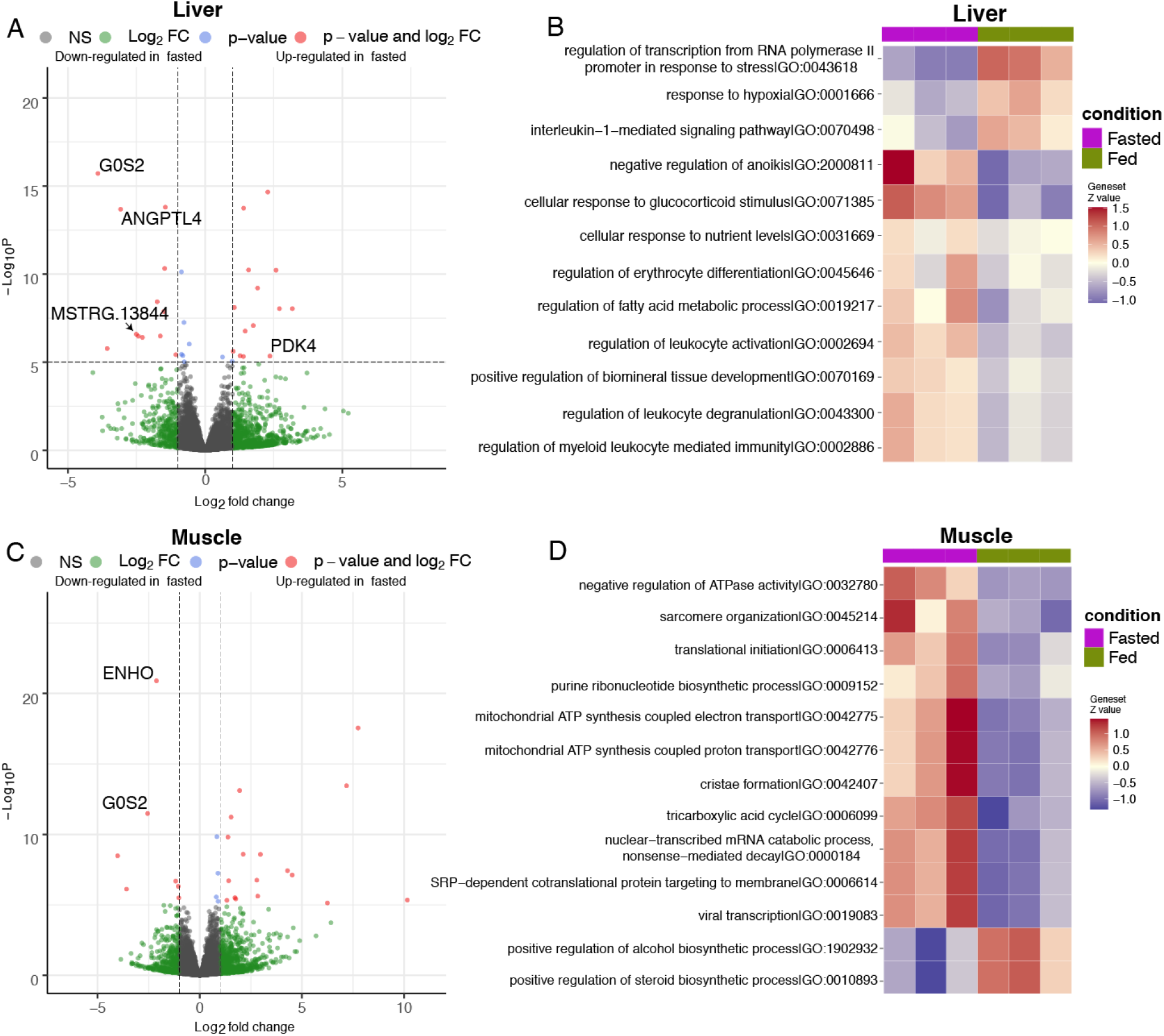
Differential Expression. **A)** Volcano plot displaying differentially expressed genes from A. Colubris liver tissue. **B**) Summary heatmap of top 12 enriched GO terms in the liver. **C)** Volcano plot displaying differentially expressed genes from A. Colubris muscle tissue. **D)** Summary heatmap of top 13 enriched GO terms in the muscle.

In the liver, the one-hour fast influenced many metabolic and homeostatic pathways **(Figure 4B),** including coenzyme biosynthetic processes, cellular responses to nutrient levels, response to hypoxia, response to carbohydrate, fatty acid metabolic processes, homeostatic processes and response to glucocorticoid stimulus **(Figure 4B).** Genes particularly affected in these processes are key regulators of metabolic flux including *PDK4, G0S2*, and *ANGPTL4*, which likely contribute to the rapid transition to lipid metabolism in the hummingbird liver during an acute food withdrawal. Induction of these genes occurs independently of PPAR signaling in the hummingbird liver as all the PPAR genes do not have any changes in expression between the fasted and fed state (**Table S8).** Therefore, in hummingbirds, the PPAR pathway does not appear to control the expression of the metabolic switch genes in the liver, at least in an acute (one hour) fast. Many newly assembled genes were also differentially expressed including *MSTRG. 13300* and *MSTRG. 13844* which we were able to functionally annotate with SwissProt as *HRG1* and *AT1B*, respectively.

The most statistically significant pathway upregulated in the fasted muscle was mitochondrial ATP synthesis coupled proton transport (GO:0042776, p=1.90E-06) **(Figure 4D).** Other key genes regulating metabolic flux were affected such as *ENHO, PPARA, G0S2* and *SREBF1* (**Figure 4C)***. ENHO* is associated with energy storage and metabolism as a precursor to the protein adropin, and was strikingly downregulated in the fasted birds (Aydin et al. 2013; Kumar et al. 2008). Interestingly, in humans, adropin is generally associated with liver and brain expression as opposed to the skeletal muscle expression we observed in *A. Colubris*. *G0S2* is the only gene that was identified as differentially expressed in both the liver and muscle tissues. While *G0S2* is known to have a significant role in liver lipid transport, a definitive role for *G0S2* within skeletal muscle has yet to be elucidated and it appears that *G0S2* is also present in mitochondria, with the speculation of several possible functions (Turnbull et al. 2016). These results point to the role of *G0S2* in hummingbird rapid metabolic flux.

## DISCUSSION

The results of our study are critical in understanding the hummingbirds’ exquisite control over rates of substrate metabolism and biosynthesis. Our positive selection analysis points to a subset of 39 genes critical to the development of nectar based life-style. Pathways such as glycolysis, the tricarboxylic acid cycle, lipogenesis and lipolysis have to function rapidly to allow for the high energetic demands of flight and reliance on nectar as the only fuel source. This was evident in the selection of genes involved in these pathways (e.g. *GAPDH, PDHA1, ACADL, HACD3*, and *BDH2*) in nectivorous bird lineages.

In our work, we look deeply into glucose and fructose uptake into the hummingbird tissues. The lack of avian GLUT4 has been previously established, but we also identified the loss of GCK. It is likely that the low levels of insulin secretion and high sustained blood glucose in hummingbirds is due in part to the lack of expression of GCK, a key regulator of insulin secretion and blood glucose homeostasis. Utilizing our assembly, annotation and expression data we speculate that hummingbird GLUT5 has transport affinity for both glucose and fructose with a higher affinity for fructose. Unlike most other animals, 50% of the hummingbirds’ diet consists of fructose (Baker, Baker, and Hodges 1998), which studies show is much more cytotoxic than glucose (Horst, Ter Horst, and Serlie 2017). As a consequence, hummingbirds rapidly sequester fructose into the muscle tissue, as evidenced by rapid declines of blood fructose levels upon fasting (Muhammad 2021). Further, we concluded that the birds are preferentially clearing fructose from circulation first and oxidizing it to CO_2_ as shown by the tracer oxidation study. Our data supports the hypothesis that both glucose and fructose are transported into the muscle cells via the GLUT5 transporter, with fructose being favored first when concentrations of both are high and glucose is imported later when fructose in blood is scarce.

Our complete assemblies of the ruby-throated hummingbird genome and transcriptome allowed for isoform level analysis of gene expression. This analysis revealed expression of a GLUT5 protein variant in the hummingbird muscle which is projected to have an internally facing active site, but due to its loss of exon three, it is unlikely that it maintains the transport activity. Future biochemical and functional studies will illuminate whether hummingbird GLUT5 is capable of fructose and glucose transport and how its variants are different from GLUT5 found in other species. We speculate that a potential biological function of this GLUT5 protein isoform is in sequestering fructose inside muscle cells rather than acting as a fructose transporter, as fructose phosphorylation capacity is low and likely cannot keep up with the rapid import. Future biochemical studies into the functionality and role of this GLUT5 protein isoform in hummingbird sugar metabolism are warranted.

We characterized the hummingbird liver and muscle expression profiles during the transition to the fasted state. These results gave insights into the drivers of rapid metabolic flux in hummingbirds. An interesting result was the upregulation of the *PDK4* gene in the fed to fasted transition. This protein kinase is located in the matrix of the mitochondria and inhibits the pyruvate dehydrogenase complex (PDH) by phosphorylating one of its subunits. Because PDH is considered the gatekeeper of the TCA cycle, its inhibition in the fasted state would shut down complete glucose oxidation and promote gluconeogenesis and fat oxidation. Expression of *PDK4* was increased 2.36 fold during fasting, results consistent with previous studies on chickens implicating glucagon as a stimulator of *PDK4* expression (Honda et al. 2017). It is possible that rapid switching from carbohydrate to fat oxidative catabolism in the fasted state is contributed to, in part, by the rapid upregulation of hummingbird *PDK4*. This suggests further study into *PDK4* molecular biology in hummingbirds. Another result calling for future biochemical and molecular studies is the downregulation of *G0S2* in both the liver and muscle tissue. *G0S2*, the G(0)/G(1) switch gene 2, is an inhibitor of Adipose triglyceride lipase (ATGL), a rate-limiting enzyme that catalyzes the first step in triglyceride hydrolysis in adipocytes. Previous studies of *G0S2* on chicken, turkey and quail have revealed avian G0S2 has 50 to 52% homology to mammalian G0S2 and suggest its importance in regulation of ATGL-mediated lipolysis (Oh et al. 2011). Our results suggest *G0S2* plays a very important role in the rapid transition of fed to fasted metabolism across multiple tissues. Lastly, we observed changes in expression of genes controlling vessel dilation and constriction, likely very important to osmoregulation (*e.g. HRG1* and *AT1B*). Hummingbird kidneys are not designed to concentrate urine as when they are feeding they must eliminate large quantities of water; however, when they are not feeding, they are susceptible to dehydration (Bakken et al. 2004). Therefore these changes in vessel dilation are likely necessary for preparing the splanchnic tissues for the osmotic shift that occurs during fasting.

In conclusion, our results have leveraged cutting-edge long and short-read sequencing technologies to generate a high quality genome assembly and annotation of the ruby-throated hummingbird. With the resources we generated, ruby-throated hummingbird genes can now be quickly cloned and expressed for further biochemical experiments, such as measuring their enzymatic properties, e.g., K_cat_ or V_max_, to compare to other avian or mammalian analogues. Expressed proteins may also be used for structural biology studies, applying either X-ray crystallography or cryoEM to generate structural maps of the proteins, then examine how the structure compares to other orthologues in dictating biological functions.

## METHODS

### Animal use and ethics statement

This study was conducted under the authority, and adheres to the requirements of, the University of Toronto Laboratory Animal Care Committee (under protocol 20011649) as well as the guidelines set by the Canadian Council on Animal Care. Twelve adult male ruby-throated hummingbirds (*Archilochus colubris*) were captured in the early summer at the University of Toronto Scarborough (UTSC) using modified box traps. The hummingbirds were individually housed in Eurocages at the UTSC vivarium on a 12h:12h light:dark cycle. The hummingbirds in these cages were provided with perches and were on an ad libitum diet of 18% weight to volume of NEKTON-Nektar-Plus (Keltern, Germany) for 2-3 months until tissue sampling occurred.

One day prior to experiment day (23 hours) 12 male birds were placed on a 33% sucrose solution ad libitum diet in place of the NEKTON-Nactar-Plus diet. Birds were then divided into a fed group (n=6) and a fasted group (n=6). One hour prior to sampling, birds from both conditions were placed in small glass jars that had perches. This restricted the birds’ ability to fly and was done in hopes of reducing energy expenditure variation between individual birds. Birds in the fed group were then provided with ad libitum 1M sucrose solution for one-hour up to sampling, which began at 10:00 h. The fasted group (n=6) was deprived of food one hour prior to sampling. The one-hour fast was chosen because previous work by Chen and Welch (2014) has shown via respirometry that this time is sufficient for the fasted hummingbird to shift from using circulating sugars to using fats for fueling metabolism.

Tissue samples were collected via terminal sampling of the hummingbirds. They were anesthetized via isoflurane inhalation and sacrificed using decapitation. Flight muscles (the pectoralis and supracoracoideus muscle) and liver were collected. Tissues were flash frozen in liquid nitrogen and subsequently stored at −80°C. In addition, one female hummingbird was also captured and sampled in the same fashion as above. This sample was used for DNA isolation for genome assembly purposes and was not subject to any experimental conditions.

### DNA Sequencing

DNA was extracted from the hummingbird from two 25mg pieces of brain tissue and two 25mg pieces of pectoralis muscle tissue with the Nanobind CBB tissue kit alpha Handbook v0.16d (4/2019) from Circulomics following the protocol for using the dounce homogenizer. DNA quality was assessed with the Thermo Scientific™ NanoDrop™ 2000/2000c Spectrophotometer. We generated a sheared nanopore library and an ultra-long nanopore library to enrich for both size and depth. For the sheared library DNA was sheared to 10kb with covaris g-tube. For the ultra-long library DNA was size-selected with the Short Read Eliminator XS Kit from Circulimics. Oxford Nanopore sequencing libraries were prepared using the Ligation Sequencing 1 D Kit (Oxford Nanopore, Oxford, UK, SQK-LSK109) according to manufacturer’s instructions and sequenced for 72 hours on 2 PromethION R9.4.1 flow cells. Nanopore reads were base-called with Guppy Software (version 3.0.6). Sequencing runs were pooled for genome assembly purposes. For shotgun Illumina sequencing, a paired-end (PE) library was prepared with the Nextera DNA Flex Library Prep Kit from Illumina and sequenced on the Illumina NovaSeq6000 (Illumina, Inc., San Diego, CA, USA). All sequencing data have been deposited at the NCBI SRA database under BioProject PRJNA811496.

### Genome assembly

The genome was assembled using both the Illumina and nanopore sequencing datasets with MaSuRCA (Zimin et al. 2013) with `FLYE_ASSEMBLY=1` and all other parameters set as default. The genome was scaffolded with RaGOO using the *C. anna* assembly (GCA_003957555.2) as a reference (Alonge et al. 2019). Assembly similarity was first checked by aligning the two assemblies with nucmer from the mummer package (Marçais et al. 2018) and assemblies were considered highly similar. Assembly completeness was checked with BUSCO using the aves lineage (Manni et al. 2021). Assembly heterozygosity was quantified with the kmer analysis toolkit (KAT) (Mapleson et al. 2017) GenomeScope (Vurture et al. 2017) and the assembly was determined to have 1.13% heterozygosity (**Figure S1)**. Repeats were annotated by first running RepeatModeler (v2.0.1) to generate a database of custom repeat annotations. The assembly was first masked with RepeatMasker (v4.0.9) using the Aves database and then further masked using the custom generated database. The genome assembly has been deposited under BioProject PRJNA811496.

### RNA extraction

RNA was extracted from approximately 40 to 50 mg of pectoralis tissue and 20 mg of liver using the Qiagen RNeasy Fibrous Tissue Mini Kit (Qiagen, Hilden, Germany). RNA quality was assessed using a nanodrop and the presence of sharp 18S and 28S rRNA on an agarose gel. RNA quality was also assessed with the Agilent 2200 TapeStation system RNA high sensitivity kit (Agilent, Santa Clara, CA) before and after polyA isolation with NEBNext^®^ Poly(A) mRNA Magnetic Isolation Module.

### RNA Sequencing

PolyA mRNA from all samples was supplemented with Spike-In RNA Variants (SRIV) set 3 from Lexogen. Libraries for Illumina sequencing were generated with NEBNext® Ultra™ RNA Library Prep Kit for Illumina and sequenced on the Illumina NovaSeq6000 (Illumina, Inc., San Diego, CA, USA). Libraries for long-read sequencing were generated with the cDNA PCR sequencing kit (SQK-PCS109) from Oxford Nanopore Technologies according to the manufacturers instructions. Libraries were each sequenced on a PromethION flow cell for 72 hours. All sequencing data have been deposited at the NCBI SRA database under BioProject PRJNA811496.

### Genome annotation and transcriptome assembly

Illumina RNA-seq reads were trimmed with trimmomatic (v0.39) with the following parameters: `SLIDINGWINDOW:4:20 LEADING:10 TRAILING:10 MINLEN:50`. Trimmed reads were then aligned to the ruby-throated hummingbird reference genome with HISAT2 (v2.2.0) (Kim et al. 2019) with the following parameters `--score-min L,0,-0.5 -k 10` to account for the high heterozygosity in the wild hummingbirds and filtered for primary alignments with Samtools (v1.9) (H. Li et al. 2009). Nanopore cDNA sequencing reads were aligned with deSALT (v1.5.4) and filtered for primary alignments with a mapping quality score greater than 50. An initial genome annotation was done by lifting over the predicted annotations from the *Calypte anna* genome annotation (GCA_003957555.2) onto our ruby-throated hummingbird assembly with LiftOff (Shumate and Salzberg 2020). These lifted over annotations were used as a reference model for hybrid transcriptome assembly with stringtie2 (Shumate et al. 2021). We ran Stringtie2 separately for each paired Illumina and nanopore sample (n=12) with the following command:

~~~
stringtie --mix {short.bam} {long.bam} -G {LiftOff_annotation.gtf}
--conservative -L -o {out.gtf} -p 10 -B -A {out.abun} -v
~~~

To filter out low evidence assembled transcripts we aligned the 12 stringtie2 gtfs with GffCompare (v0.11.2) (Pertea and Pertea 2020). The 12 gtf files were merged with stringtie merge and filtered to retain transcripts that had evidence from at least two of the 12 gtfs. Gene and transcript abundance measurements were computed against the final merged and filtered gtf file with the same command as above and the addition of the -e flag:

~~~
stringtie --mix {short.bam} {long.bam} -G {filtered_merged.gtf}
--conservative -L -o {out.gtf} -p 10 -B -A -e {out.abun} -v
~~~

To correct for transcripts assigned to the incorrect gene locus during stringtie’s merge function we ran the R package IsoformSwitchAnalyzeR (Vitting-Seerup and Sandelin 2019) with fixStringTieAnnotationProblem = TRUE. We generated the transcript count matrix files using the prepDE.py3 script from stringtie2 and the gene count matrix files were generated using the abundance measurements from the gtf and the gene to transcript associations from the IsoformSwitchAnalyzeR output. The ruby-throated hummingbird transcriptome assembly is available on zenodo (DOI:10.5281/zenodo.6363333). Protein predictions from the transcriptome were done with TransDecoder (v5.5.0) (https://github.com/TransDecoder/TransDecoder). Genes that were not annotated in the anna’s hummingbird reference were first confirmed to have functional open reading frames by identifying a corresponding protein prediction from the TransDecoder output. They were then functionally annotated by Blast (v2.2.31+)(Camacho et al. 2009) to the Swiss-Prot database (Boeckmann 2003) and run through the InterProScan5 (v5.44-79.0) pipeline (Jones et al. 2014).

### Differential expression

Differential gene expression was done with DESeq2 (Love, Huber, and Anders 2014) filtering for genes with at least 10X coverage in at least four of the six samples per tissue. The three fasted samples and three fed samples were compared separately for the liver and muscle tissue and significantly differentially expressed genes were determined using adjusted p-values beneath 0.1. Isoform level expression was quantified with Ballgown (Frazee et al. 2015) and isoform level FPKM values were compared across the muscle and liver tissues. Significantly upregulated pathways were determined with the GeneTonic R package (Marini et al. 2021) for both liver and muscle.

### Gene loss analysis

We did not identify the GCK gene in our transcriptome annotation or in the functional annotation of the predicted proteins. As further validation we used Blast (v2.2.31+) using the chimney swift (*Chaetura pelagica*) and chicken (*Gallus gallus*) GCK gene sequence and protein sequence to both the ruby-throated hummingbird predicted protein set and genome. The Blast search did not uncover any hits that we could determine to be open reading frames. We then aligned the chicken reference genome (GCA_000002315.5) to the ruby-throated hummingbird genome with minimap2 (H. Li 2016) with the following parameters: `minimap2 -x asm20 -c --eqx`. We noted that the region with the GCK gene in the chicken genome is non-syntenic to any of the ruby-throated hummingbird DNA sequence. The paf output file was processed with rustybam (https://github.com/mrvollger/rustybam) and plotted with SafFire (https://mrvollger.github.io/SafFire/). To further ensure that there was no expression of the GCK gene in the ruby-throated hummingbird we mapped all the RNA-seq reads to the chimney swift GCK gene sequence with Bowtie2 (Langmead and Salzberg 2012) allowing for multiple mismatches by using the very-sensitive-local flag. None of the RNA-seq reads mapped to the chimney swift GCK sequence. Lastly, we also validated that the GCK gene was not present in the annotation of Anna’s hummingbird either.

### Positive selection analysis

For molecular evolution analyses, we used a consensus tree topology based on molecular phylogenies generated by Hacket et al., Oliveros et al., and Prum et al (Hackett et al. 2008; Oliveros et al. 2019; Prum et al. 2015). Species were chosen to give outgroups in multiple clades, as well as provide species as a sister lineage for each nectivorous lineage, where such a species existed in the publicly available genome databases. Additionally, species were added that evolved between nectivorous lineages to highlight the convergent nature of the phenotype. Additionally, adding lineages between nectivorous lineages allowed us to ensure that the branches we tested for positive selection to the best of our ability matched the branches where the transition to nectivory happened in more species rich phylogenies. Proteomes for all species selected were downloaded from NCBI and clustered with cd-hit (v4.8.1) with a sequence identity threshold of 98% to remove redundancy in the datasets (W. Li and Godzik 2006). Orthologous gene groups were generated by running the clustered proteomes through the OrthoFinder (v2.3.12) pipeline (Emms and Kelly 2019). 1-to-1 orthology groups were determined by selecting all single copy genes that were contained in all species.

For each 1-to-1 orthology group (OG), the branch-site test of positive selection was performed using codeml in PAML v4.10 (https://github.com/abacus-gene/paml; Yang 2007) to detect genes under positive selection in nectivorous bird lineages. Using the phylogenetic tree reported in phylogenies generated by Hacket et al., Oliveros et al., and Prum et al (Hackett et al. 2008; Oliveros et al. 2019; Prum et al. 2015), the topology was unrooted using the ete3 toolkit (Huerta-Cepas, Serra, and Bork 2016), and foreground branches were assigned to the following nectivorous lineages: *Grantiella picta, Promerops cafer, Leptocoma aspasia*, and the clade of *Calypte anna + Archilochus colubris*. A likelihood ratio test (LRT) was performed for each OG, with the branch-site model A (specified in Yang 2007) as the alternative model and model A with a fixed ω = 1 as the null model. LRT statistics were converted to p-values using pchisq in R v.3.5.0 (R Core Team 2018). To provide a conservative estimate of genes under positive selection among nectivorous lineages, each OG with a statistically significant LRT (p ≤ 0.05) was also required to possess at least one site under positive selection with a posterior probability ≥ 0.95 (according to the Bayes empirical Bayes analysis of positive selection included in the codeml branch-site test of positive selection). Genes under positive selection are reported in **Table S3**. Intermediate files from this analysis are available on zenodo (DOI:10.5281/zenodo.6363333).

### Protein structure models

Structures for full length GLUT5 (isoform X1) and for the alternative spliced variant (isoform X2) where modeled with AlphaFold v2.01 (https://github.com/deepmind/alphafold) using default settings without templates to avoid model bias (Jumper et al. 2021). A reduced version of the BFD database (https://bfd.mmseqs.com/), optimized for speed and lower hardware requirements, was employed during multi-sequence alignment (MSA). The overall confidence measure (predicted local-distance difference test, pLDDT) for the generated models was > 75, which generally indicates good backbone prediction. Atomic models with highest scores at the overall confidence measure were selected (86.2 for isoform X1 and 84.5 for isoform X2). pLDDT is a per-residue confidence metric, and as such, can be used to monitor how the model confidence varies along the chain. Very low confidence regions (<50) included flexible amino- and C-terminal ends that were removed from the models.

### Tracer oxidation study

Four ruby-throated hummingbirds were fasted for 1 hour, following which they were placed in a 500 ml respirometry container and baseline fasting breath delta ^13^C breath stable isotope signature and respiratory exchange ratio (RER) recording (see (Dick et al. 2020) for respirometry and breath stable isotope set up). After 5 min the birds were then fed a 150 ml of a 20% sucrose solution with sucrose enriched with ^13^C on all six carbons of the glucose (sucrose (glucose-^13^C6, 98%), Cambridge Isotope Laboratories, Tewksbury, MA, USA] or fructose [d-sucrose (fructose-^13^C6, 98%), Cambridge Isotope Laboratories] portion of the sucrose molecule. The birds were fed through a 1 ml syringe in the lid of the respirometry jar which allowed for continuous breath measurements, and previous training allowed for quick consumption of the sucrose solutions. The time of feeding was recorded and used as t=0. The respiratory measurements continued over the next 20 minutes to measure the rise and start of the fall of RER, representing the switch from fasted to fed. The birds were then returned to their cages and repeated the process again 1 week later with the other sucrose solution, with 2 birds starting with fructose-enriched, and 2 birds starting with the glucose-enriched. RER was analyzed following (Dick et al. 2020), and tracer oxidation rate analyzed following (M. D. McCue et al. 2010), and were averaged for each minute over the course of 20 minutes. The time to and peak tracer oxidation rate was analyzed using a pair t-test.

## SUPPLEMENTAL FIGURES

**Figure S1.**
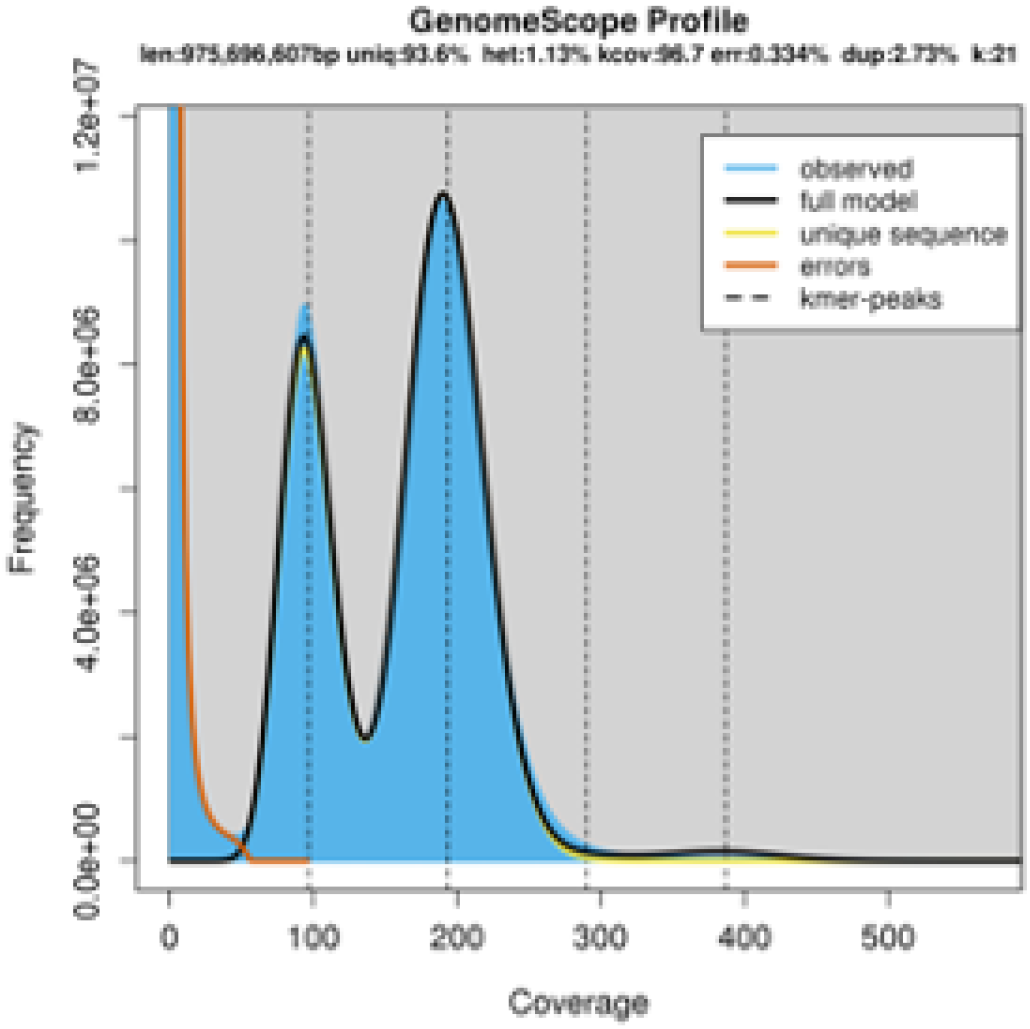
Genome-scope plot. Coverage and kmer frequency plot using the Illumina gDNA reads and the MaSURcA assembly of the ruby-throated hummingbird.

**Figure S2.**
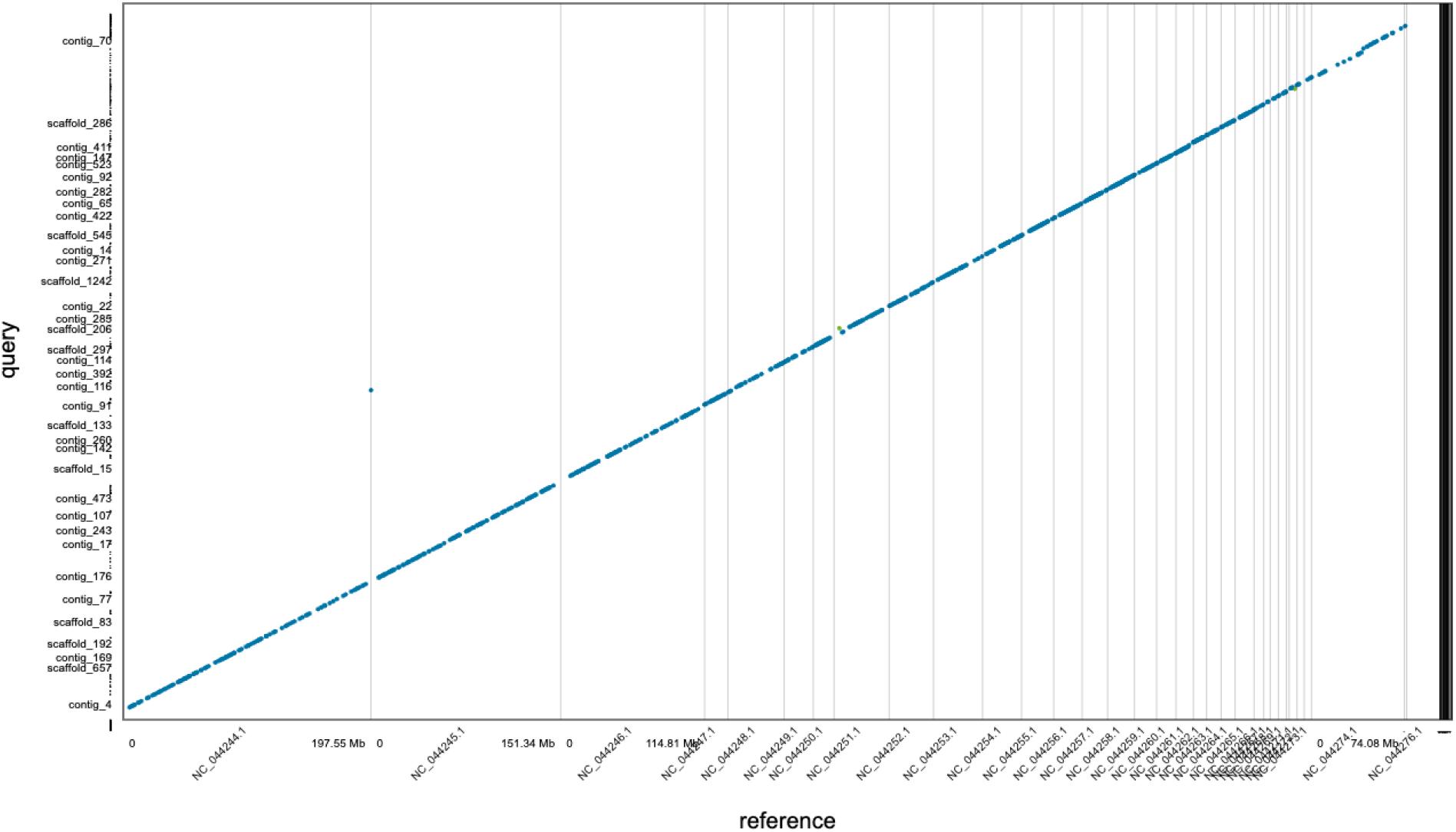
Assembly alignment. Whole genome alignment of the unscaffolded ruby-throated hummingbird MaSURcA assembly to the Anna’s hummingbird assembly (GCA_003957555.2).

**Figure S3.**
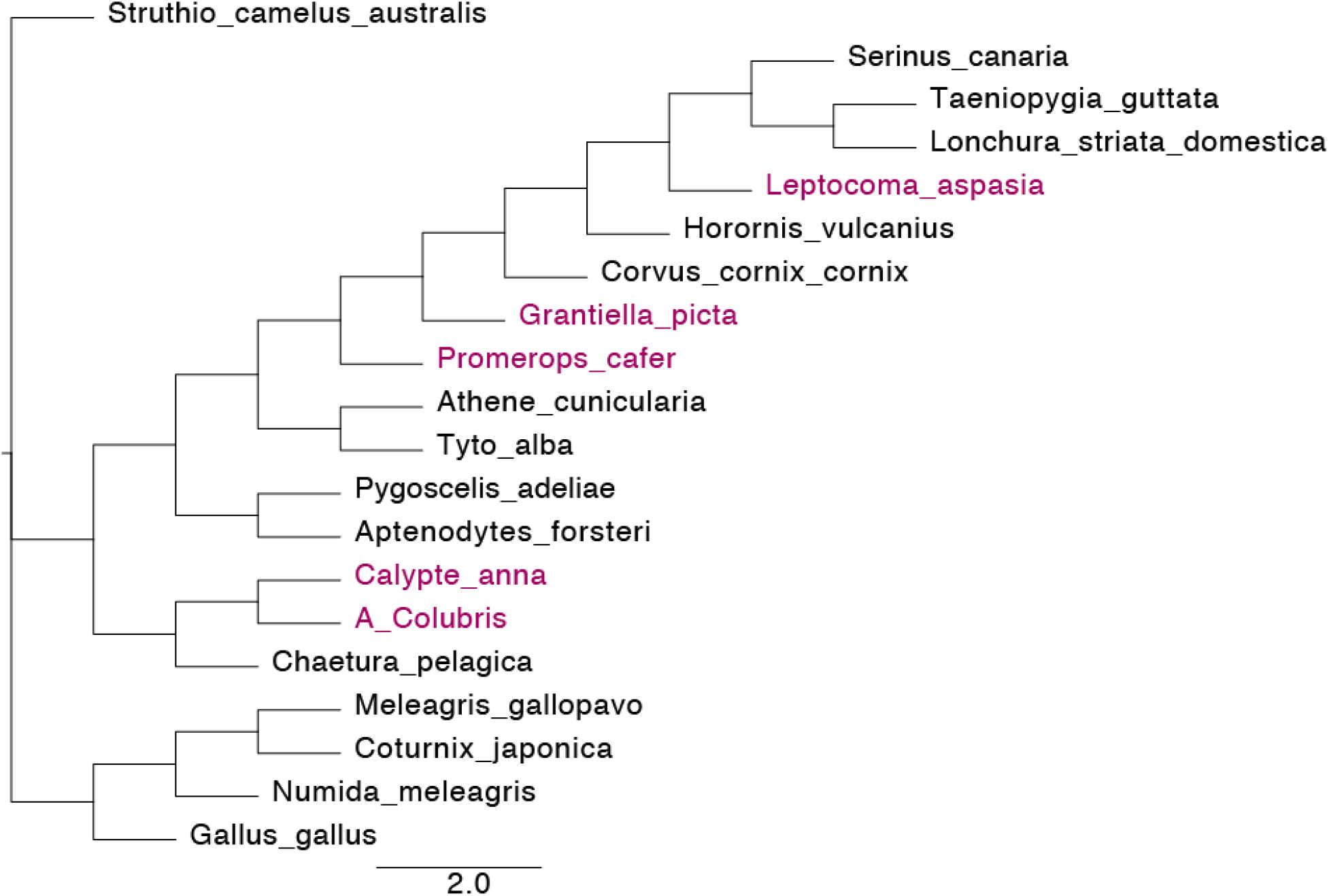
Phylogenetic tree. Bird phylogeny for positive selection analysis. The nectivorous branches of interest, known as the foreground branches, are written in pink, background branches in black.

**Figure S4.**
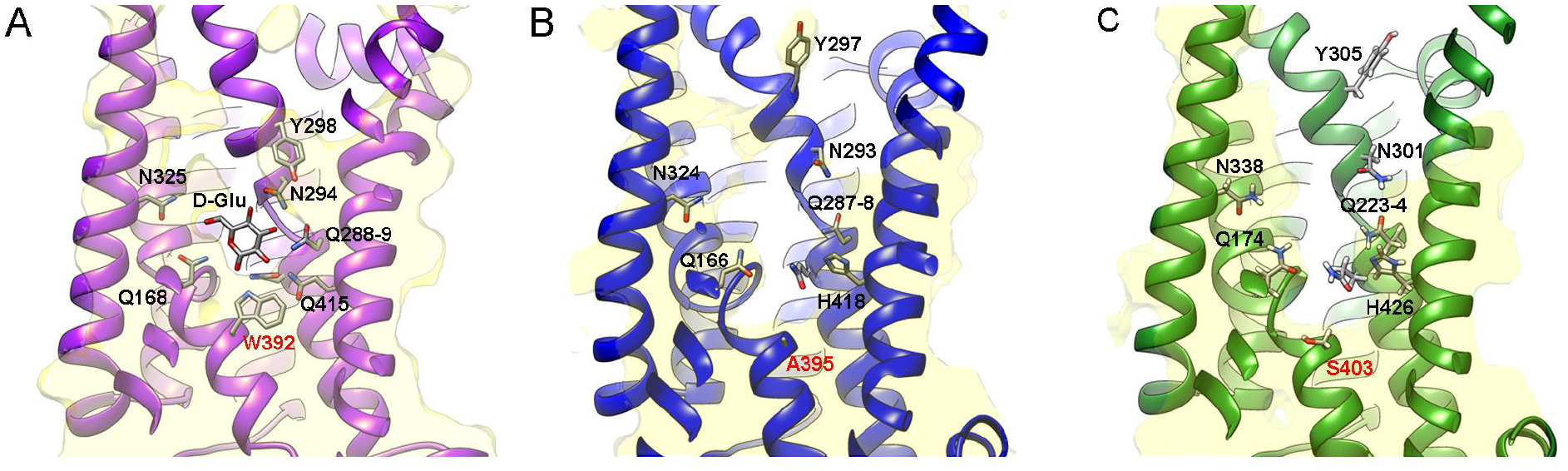
Sugar binding site in GLUT structures. Ribbon representations of the sugar binding pocket of atomic structures for: **A)** bacterial XylE in complex with D-Glucose (pdb code 4bgz, Sun et al., 2012), **B)** rat GLUT5 (pdb code 4ybq, Nomura et al., 2015), and **C)** an atomic model for ruby-throated hummingbird GLUT5 generated with AlphaFold. Residues involved in hydrogen bonding interactions with the glucose in XylE are labeled on allthe three structures. The semitransparent overlays in yellow depict central sections of the volumes for the atomic models and show the changes at the bottom of the binding pocket where different amino acids are found in GLUT transporters (residues labeled in red).

**Figure S5.**
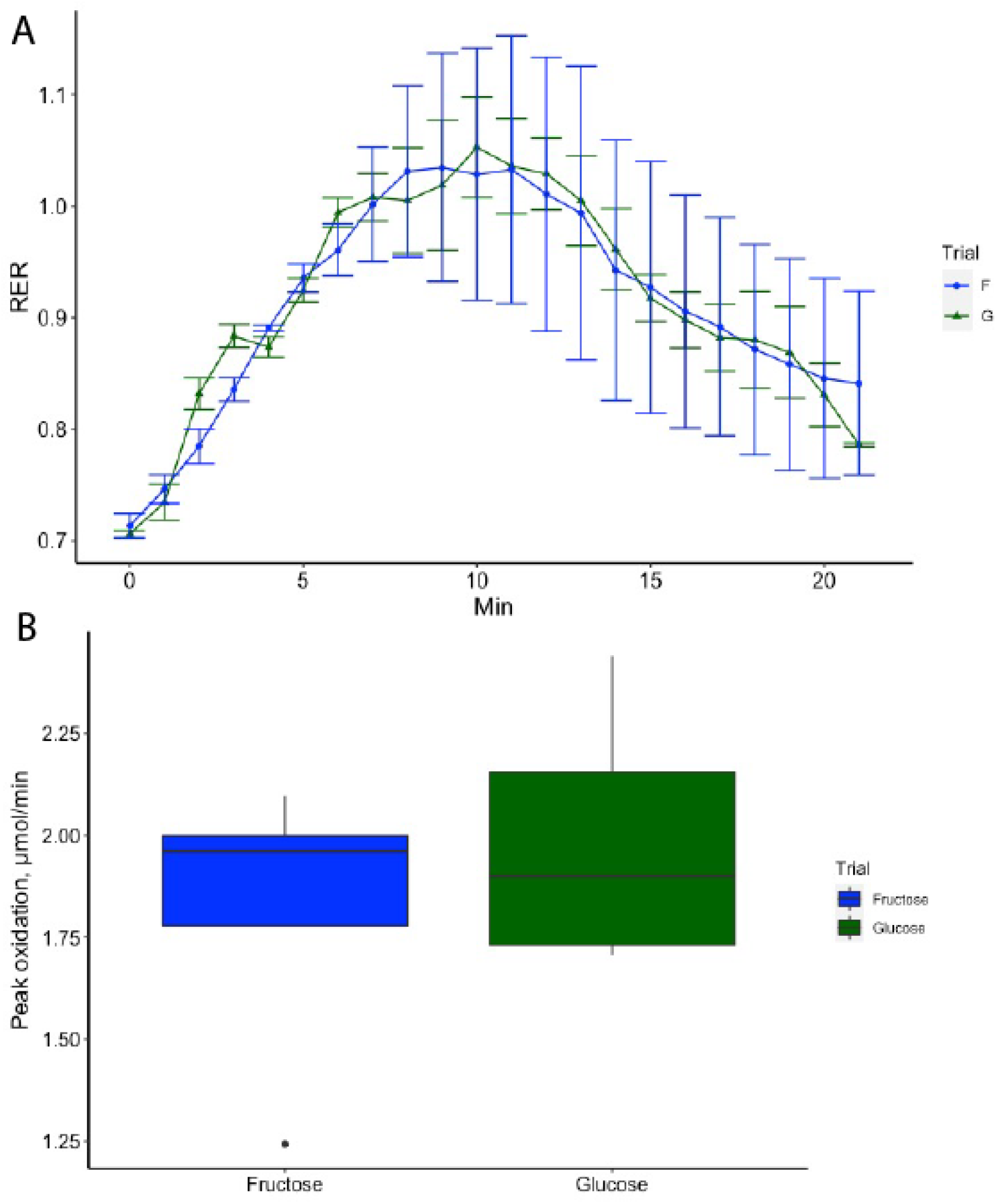
Tracer oxidation. **A)** respiratory exchange ratio 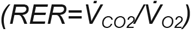 (RER) of glucose and fructose over a twenty minute time-course. **B)** Peak tracer oxidation between the enriched sucrose solutions (p = 0.66, paired t-test)

**Figure S6.**
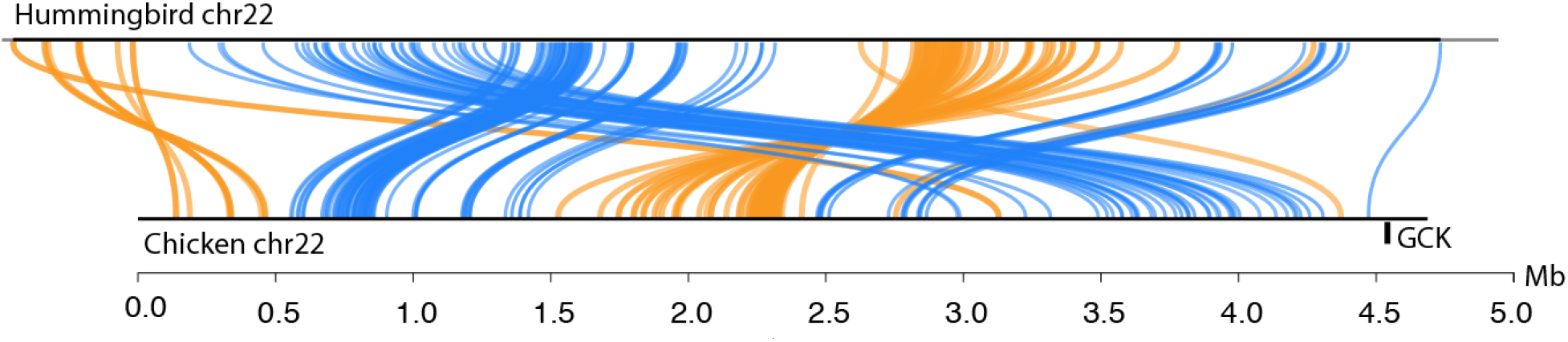
Chicken and hummingbird synteny. Synteny plot showing syntenic sequence between the chicken and ruby-throated hummingbird in the forward direction (blue) and syntenic sequence that is reserved (orange).

## Supplementary table legends

**Table S1.** *A. Colubris Genome BUSCO results*.

**Table S2.** *A. Colubris Genome repeat annotations*.

**Table S3.** *Positively selected genes in nectivory with Panther gene function IDs*.

**Table S4.** *Differentially expressed genes in the fasted versus fed A. Colubris liver*.

**Table S5.** *Differentially expressed genes in the fasted versus fed A. Colubris muscle*.

**Table S6.** *Gene ontology pathway analysis of differentially expressed genes in the A. Colubris liver*.

**Table S7.** *Gene ontology pathway analysis of differentially expressed genes in the A. Colubris muscle*.

**Table S8.** *Gene expression results from the genes involved in the PPAR signaling pathway in both the fasted and fed muscle and liver*.

